# Complementary Insights from Environmental DNA and Environmental RNA Metabarcoding for Marine Biodiversity Assessment Around San Andrés Island, Colombia

**DOI:** 10.64898/2026.06.03.730006

**Authors:** Sean K Bedingfield, Caroline Vanegas Moreno, Alexander F More

## Abstract

Environmental DNA (eDNA) metabarcoding has become a cornerstone of marine biodiversity monitoring, yet it recovers genetic material irrespective of organism viability and may therefore conflate historical and contemporary community signals. Environmental RNA (eRNA), derived from less stable ribonucleic acid, is hypothesized to be biased toward metabolically active organisms and may provide a more temporally resolved snapshot of living communities. Here we present a paired eDNA/eRNA metabarcoding comparison across a tropical marine seascape, analyzing 19 co-sampled sites spanning coral reefs, mangroves, a seagrass bed, shipwrecks, a cenote, and coastal infrastructure around San Andrés Island, Colombia. To our knowledge this is the first in situ, ecosystem-scale paired eDNA/eRNA survey of the broad eukaryotic community across multiple natural habitat types in a tropical marine system, extending mesocosm and freshwater work (e.g., Giroux et al., 2022) to a field setting. Using COI-region amplicon sequencing processed by NatureMetrics, we recovered 1,944 operational taxonomic units (OTUs) across the 19 paired sites. Of these, 1,015 (52.2%) were detected by both approaches, 305 (15.7%) were unique to eDNA, and 624 (32.1%) were unique to eRNA. The eRNA-unique fraction was taxonomically enriched for groups including diatoms (class Bacillariophyceae, phylum Ochrophyta), ciliates, and other protists. Paired Wilcoxon signed-rank tests showed that eRNA recovered significantly higher OTU richness (median 239 vs. 207; W = 36, p = 0.016) and Shannon diversity (median 3.64 vs. 3.38; W = 40, p = 0.026) than eDNA. The mean per-site Jaccard similarity between paired samples was 0.40, indicating substantial turnover in the rare-taxon composition recovered by each method. Principal coordinates analysis of Bray-Curtis dissimilarity showed that habitat type structured abundance-weighted community composition (PERMANOVA F = 2.49, p = 0.001) whereas molecular method did not (F = 1.37, p = 0.107). A PERMDISP test found homogeneous multivariate dispersion between methods (F = 0.01, p = 0.92), reinforcing the absence of a method effect, but significant dispersion heterogeneity among habitats (F = 24.0, p < 0.01), so the habitat result is interpreted with caution. Indicator species analysis identified 73 OTUs significantly associated with one template: eDNA indicators were dominated by dinoflagellates (Dinophyceae) and eRNA indicators by diatoms (Bacillariophyceae) and fungi, consistent with an eRNA bias toward metabolically active microbial eukaryotes. A read-weighted overlap analysis showed that although eRNA-unique OTUs outnumbered eDNA-unique OTUs roughly two to one, the large majority of reads (>95%) fell in shared OTUs, so method-unique detections are predominantly rare taxa. We discuss the complementary value of eRNA for marine monitoring, with the seagrass habitat — where eRNA reduced masking by terrestrial plant material — as the clearest use case, and propose, rather than prescribe, the integration of eRNA into routine programs.

## 1. Introduction

Environmental DNA (eDNA) metabarcoding has transformed aquatic biodiversity assessment over the past decade, enabling non-invasive, cost-effective surveys of whole communities from water samples. By sequencing extracellular genetic material shed by organisms, eDNA approaches can detect hundreds of taxa simultaneously without visual surveys or physical capture, and have been adopted worldwide for fisheries management, invasive-species detection, and ecosystem-health monitoring (Thomsen & Willerslev, 2015; Miya, 2022).

However, eDNA carries a well-recognized limitation: DNA molecules can persist in the water column for days to weeks after an organism has died or left a location (Harrison et al., 2019). eDNA signals therefore reflect a temporal and spatial average of community presence rather than a strictly contemporaneous snapshot, and in dynamic marine settings advection can transport genetic material across habitat boundaries.

Environmental RNA (eRNA) offers a theoretically complementary signal. Because RNA degrades more rapidly than DNA in the extracellular environment, eRNA-based detections are hypothesized to be more strongly weighted toward metabolically active organisms near the time and place of sampling (Pochon et al., 2017; Wood et al., 2020). Paired eDNA/eRNA approaches have been explored in soil and freshwater systems and in marine sediment mesocosm and estuarine work (Laroche et al., 2017; Giroux et al., 2022), but in situ applications across a natural tropical marine seascape remain rare. Our study builds on this foundation rather than claiming priority over it: we extend the paired eDNA/eRNA concept to an ecosystem-scale field survey spanning multiple natural habitat types.

The Seaflower Biosphere Reserve surrounding San Andrés Island, Colombia, is a UNESCO-designated reserve harboring exceptional marine biodiversity across coral reefs, mangroves, seagrass meadows, and anthropogenic structures such as shipwrecks and submarine infrastructure (Sánchez et al., 2019, 2021), providing a natural mosaic in which to evaluate eRNA as a complement to eDNA.

In this study we present a paired eDNA/eRNA sampling campaign across 19 co-sampled sites around San Andrés Island (Figure 1). Our objectives were to: (1) compare the taxonomic composition and diversity recovered by each template, using paired statistical tests; (2) determine whether molecular method or habitat type drives community composition, using ordination, PERMANOVA, and PERMDISP; (3) identify taxa preferentially associated with eDNA or eRNA using indicator species analysis; (4) place presence/absence overlap in an abundance context using read-weighted overlap and rarefaction; and (5) assess the feasibility of incorporating eRNA into routine marine biodiversity monitoring.

**Figure 1.**
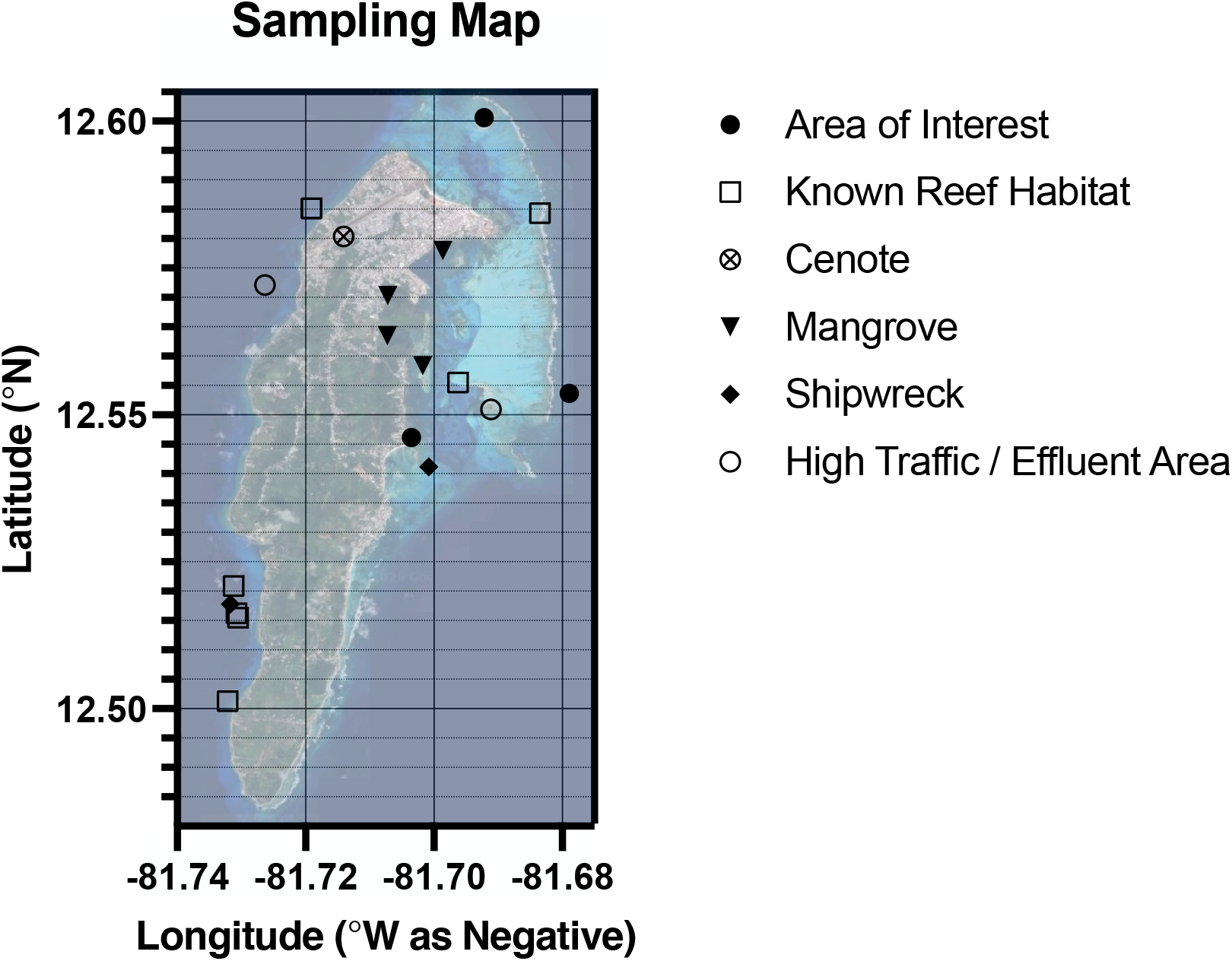
Sampling sites around San Andrés Island, Colombia, within the Seaflower Biosphere Reserve, categorized by habitat type. Of twenty sites sampled, 19 yielded paired eDNA/eRNA data.

## 2. Materials and Methods

### 2.1 Study Area

Sampling was conducted around San Andrés Island (approximately 12.58°N, 81.70°W), within the Seaflower Biosphere Reserve (≈180,000 km^2^). Twenty sites were sampled across six habitat categories: coral reef (n = 12), mangrove (n = 3), seagrass (n = 1), shipwreck (n = 2), cenote (n = 1), and coastal infrastructure (n = 1). One mangrove site (Old Point) yielded eDNA data only and is excluded from all paired analyses; the 19 sites with paired eDNA/eRNA data form the basis of this study. Two sites (Plaza de Toros and The Lighthouse) were sampled as field duplicates rather than independent biological replicates, and are described as such throughout. Site coordinates are taken from the field sample manifest and shown in Figure 1; per-site latitude and longitude are tabulated in the Supplementary Material.

### 2.2 Sample Collection

Water samples were collected using the EQO Osprey platform, an automated in situ filtration system. A target volume of seawater was filtered per sample onto membrane filters, which were preserved in situ with a proprietary stabilization buffer (EQO) intended to co-preserve DNA and RNA. Samples were maintained under cold-chain conditions during transport to the laboratory.

### 2.3 Nucleic Acid Extraction and Reverse Transcription

Preserved filters were processed by EQO. DNA was extracted using a magnetic-bead environmental DNA protocol (Mag-Bind Environmental DNA Kit, Omega Bio-Tek; cf. Spens et al., 2017) incorporating proteinase K lysis and an inhibition-removal step, and extract concentration was quantified using a Qubit dsDNA assay (broad-range chemistry; Thermo Fisher Scientific).

For the eRNA fraction, RNA was DNase-treated and reverse-transcribed to complementary DNA (cDNA) prior to amplification, with a no-reverse-transcriptase control to confirm RNA-template specificity. The cDNA was then carried through the same amplicon workflow as the DNA fraction.

### 2.4 Metabarcoding and Bioinformatics

COI-region amplicon libraries were prepared from both eDNA and cDNA templates using an identical primer set and sequenced on an Illumina platform (NatureMetrics, Oxford, UK). Reads were demultiplexed and paired reads merged (USEARCH; Edgar, 2010); primers were trimmed with cutadapt (Martin, 2011); reads were quality-filtered by expected error; and sequence variants were denoised with the UNOISE algorithm (Edgar, 2016). Variants were clustered into OTUs at 97% similarity (UPARSE; Edgar, 2013) with chimera removal and a low-abundance read filter. Taxonomy was assigned by BLAST (Altschul et al., 1990; Camacho et al., 2009) against the NCBI nucleotide database with stringent identity and coverage thresholds, reconciled to the GBIF backbone taxonomy (via rgbif; Chamberlain et al., 2023) using tiered percent-identity cutoffs for species-, genus-, and higher-rank assignments. Unassigned OTUs and human/domesticated-animal sequences were removed prior to analysis.

### 2.5 Statistical Analyses

All paired analyses used the 19 sites with both eDNA and eRNA data. For each site we calculated OTU richness, Shannon diversity (H′ = −Σ p_i_ ln p_i_), Jaccard similarity (J = |A ∩ B| / |A ∪ B|), and phylum- and class-level relative read abundance. A read-weighted overlap analysis quantified the total reads in shared versus method-unique OTU categories, complementing the presence/absence (Venn) view.

#### 2.5.1 Paired diversity tests

Because eDNA and eRNA samples are paired by site, differences in OTU richness and Shannon diversity between templates were tested with two-sided Wilcoxon signed-rank tests across the 19 sites.

#### 2.5.2 Rarefaction

To control for variation in sequencing depth (eDNA: 16,758–106,498 reads; eRNA: 21,144– 184,632 reads), samples were rarefied by subsampling without replacement to the minimum observed depth (16,758 reads), and rarefaction curves were generated to assess saturation. Rarefaction is used here as a supporting robustness check rather than as definitive proof; the choice of normalization for metabarcoding data remains debated.

#### 2.5.3 Multivariate analysis, PERMANOVA, and PERMDISP

Community composition was compared using Bray-Curtis dissimilarity on the rarefied OTU abundance matrix and ordinated by Principal Coordinates Analysis (PCoA). PERMANOVA (Anderson, 2001; 999 permutations) tested whether composition differed by molecular method and by habitat type, with separate within-method habitat tests. Because PERMANOVA can be sensitive to within-group dispersion differences, we additionally ran a test of multivariate homogeneity of group dispersions (PERMDISP; Anderson, 2006) for both groupings, so that location and dispersion effects could be distinguished.

#### 2.5.4 Indicator species analysis

To identify taxa preferentially recovered by each template, we applied indicator value analysis (IndVal; Dufrêne & Legendre, 1997) to OTUs grouped by method, assessing significance with 999 permutations and correcting for multiple testing with the Benjamini-Hochberg false-discovery-rate procedure. OTUs with an indicator value above 0.3 and an FDR-adjusted q < 0.05 were retained.

## 3. Results

### 3.1 Sampling Sites

Nineteen sites yielded paired eDNA/eRNA data across six habitat types (Figure 1). Coral reef sites predominated (n = 12), with the remaining habitats — mangrove, seagrass, shipwreck, cenote, and coastal infrastructure — represented by one or two sites each, a sampling imbalance that is relevant to interpretation of the habitat-level analyses below.

### 3.2 Overall Detection and Overlap

Across the 19 paired sites we recovered 1,944 OTUs. Of these, 1,320 were detected by eDNA and 1,639 by eRNA. 1,015 OTUs (52.2%) were shared, 305 (15.7%) were eDNA-unique, and 624 (32.1%) were eRNA-unique (Figure 2). To ensure the method comparison is not biased by an unpaired sample, these totals are restricted to the 19 sites with both templates; the eDNA-only Old Point site is excluded here and reported separately in the Supplementary Material, where its inclusion is shown not to affect any conclusion.

**Figure 2.**
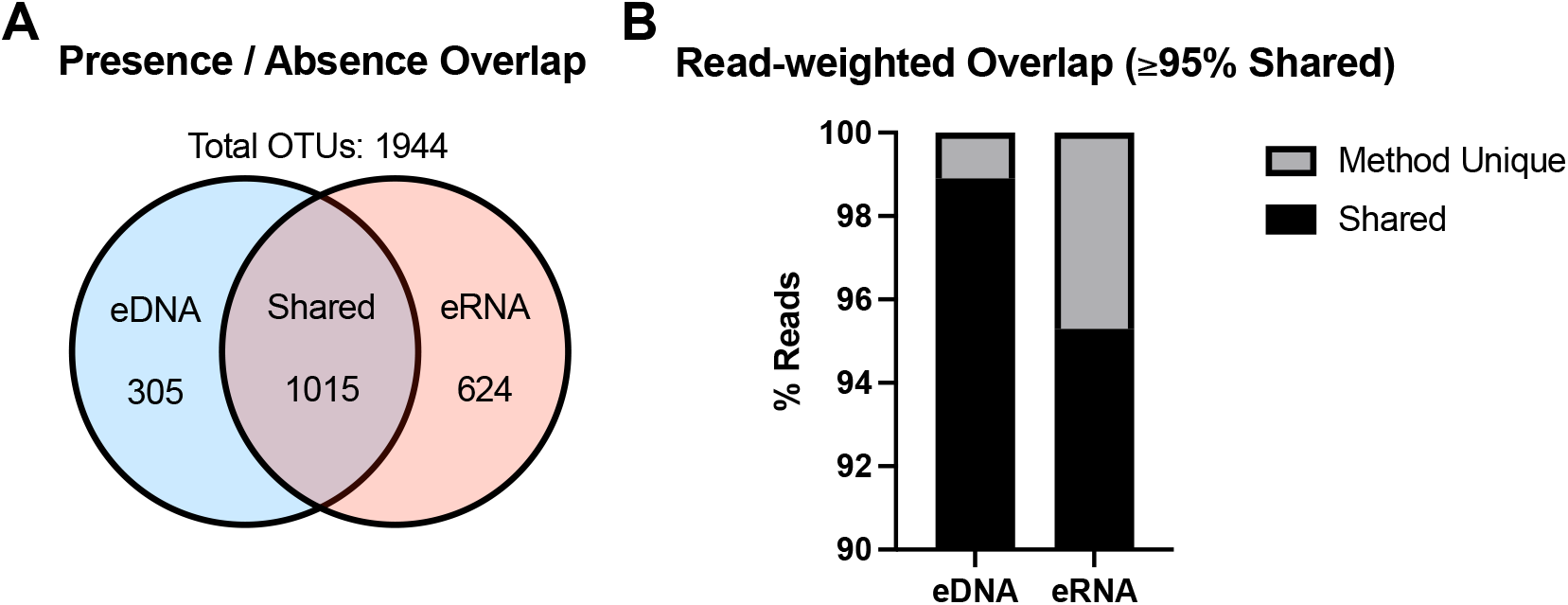
(A) Global OTU overlap between eDNA and eRNA (presence/absence, 19 paired sites). (B) Read-weighted overlap: total reads in shared versus method-unique categories.

### 3.3 Site-Level Richness and Rarefaction

OTU richness varied substantially across sites (Figure 3). eRNA recovered higher OTU richness than eDNA at 14 of 19 sites, and a paired Wilcoxon signed-rank test confirmed the difference was significant (W = 36, p = 0.016; median 239 for eRNA vs. 207 for eDNA). Rarefaction to the minimum observed depth (16,758 reads) left this pattern essentially unchanged, indicating it is not an artifact of differential sequencing effort; the observed-versus-rarefied richness comparison and rarefaction curves are provided in the Supplementary Material.

**Figure 3.**
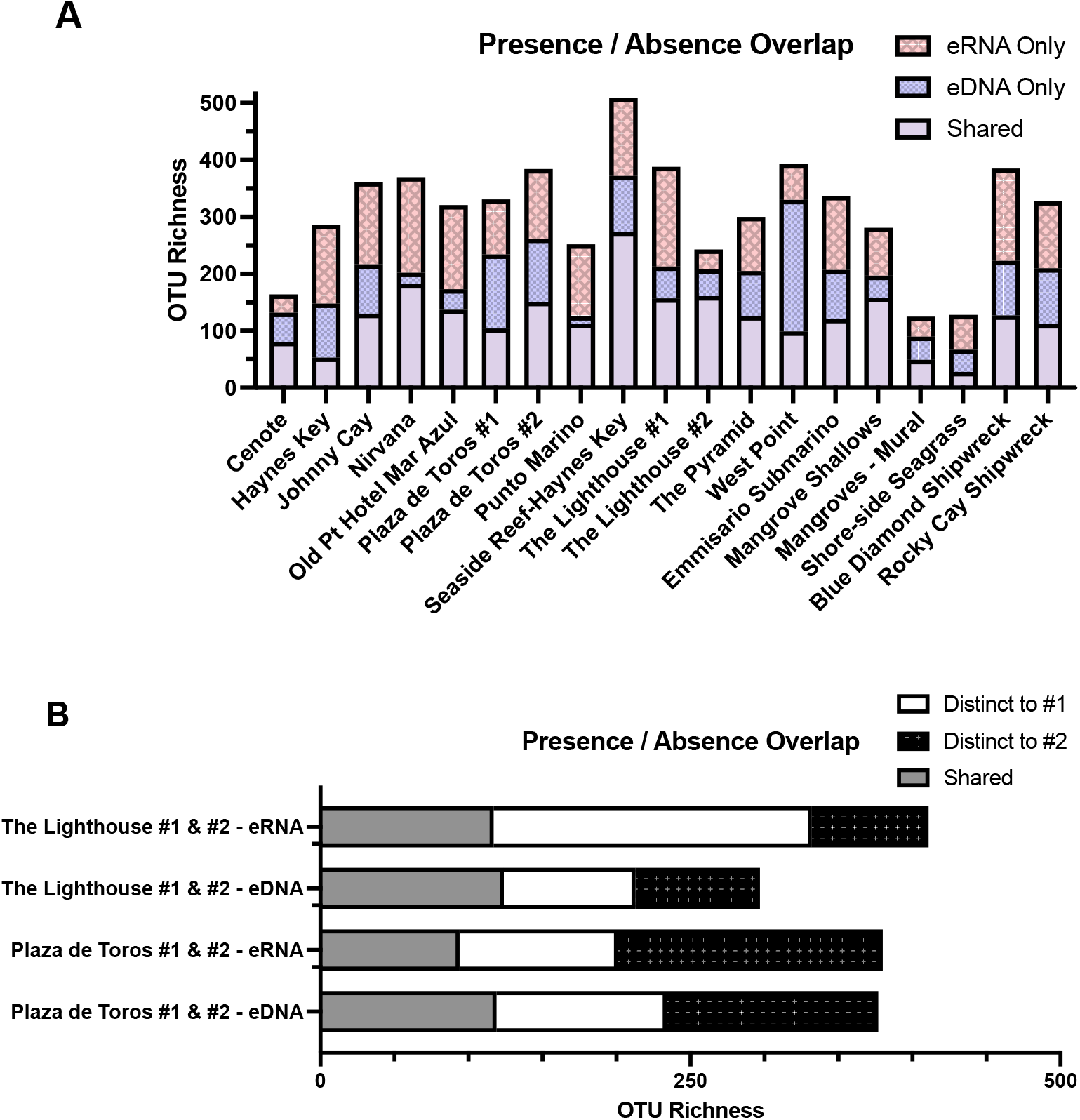
(a) OTU richness by site, with each bar partitioned into shared, eDNA-only, and eRNA-only OTUs and the counts annotated directly on the bars. Rarefied richness is provided in the Supplementary Material. (b) OUT richness compared within method between duplicates at two sites.

**Figure 4.**
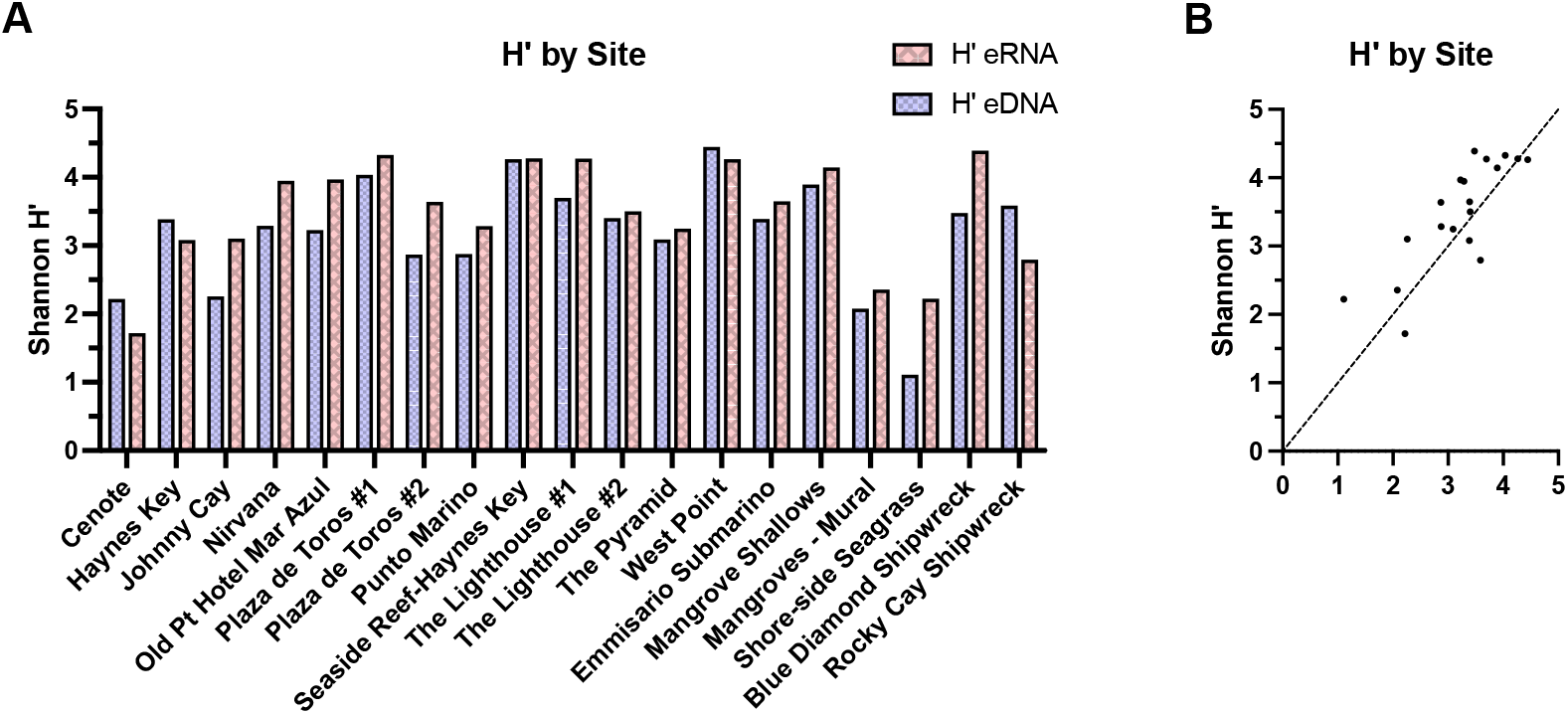
(A) Per-site Shannon diversity for eDNA and eRNA. (B) eDNA-versus-eRNA scatter with the 1:1 line; points above the line indicate higher eRNA diversity.

The five sites where eDNA recovered higher richness than eRNA (Plaza de Toros #1, West Point, Cenote, Mangroves–Mural, and The Lighthouse #2) did not share a common habitat or obvious environmental characteristic; they span coral reef, cenote, and mangrove settings. We therefore interpret these as local, likely stochastic departures from the general pattern rather than evidence of a systematic habitat-linked reversal.

### 3.4 Read-Weighted Overlap

Although the presence/absence view shows eRNA detecting roughly twice as many unique OTUs as eDNA, the read-weighted analysis shows that more than 95% of reads fall in shared OTUs (Figure 2B). eDNA-unique OTUs contributed only 14,261 reads (1.1% of eDNA reads), and eRNA-unique OTUs 70,041 reads (4.7% of eRNA reads). Method-unique detections are therefore predominantly low-abundance, rare taxa rather than dominant community members.

### 3.5 Diversity Comparison

Shannon diversity (H′) was higher for eRNA at 15 of 19 sites, and a paired Wilcoxon signed-rank test confirmed the difference was significant (W = 40, p = 0.026; median 3.64 for eRNA vs. 3.38 for eDNA). The largest single-site difference occurred at the Shore-side Seagrass site (eRNA H′ = 2.222 vs. eDNA H′ = 1.110), discussed below in the context of terrestrial-input masking.

We report per-site diversity for each template separately rather than a single pooled-dataset diversity value. Pooling eDNA and eRNA reads per OTU and recomputing H′ can yield a value lower than one of the contributing templates when the two differ markedly in evenness, because pooling reweights relative abundances; a pooled metric is therefore not a straightforward measure of added information. To avoid this ambiguity, the pooled-diversity computation and its behavior are documented in the Supplementary Material rather than presented as a primary result.

### 3.6 Taxonomic Composition

At the phylum level, eDNA read abundance was dominated by Arthropoda (51.1%), whereas eRNA showed elevated Annelida (13.9% vs. 4.5% in eDNA) and Ochrophyta (8.9% vs. 4.8%) (Figure 5A). Because COI recovers a very broad range of taxa, we also summarize composition at class level (Figure 5B), which is more informative for several groups: the eRNA enrichment in Ochrophyta is driven largely by diatoms (class Bacillariophyceae: 8.2% of eRNA vs. 3.7% of eDNA reads), while dinoflagellates (class Dinophyceae) are comparatively enriched in eDNA (9.5% vs. 2.9%).

**Figure 5.**
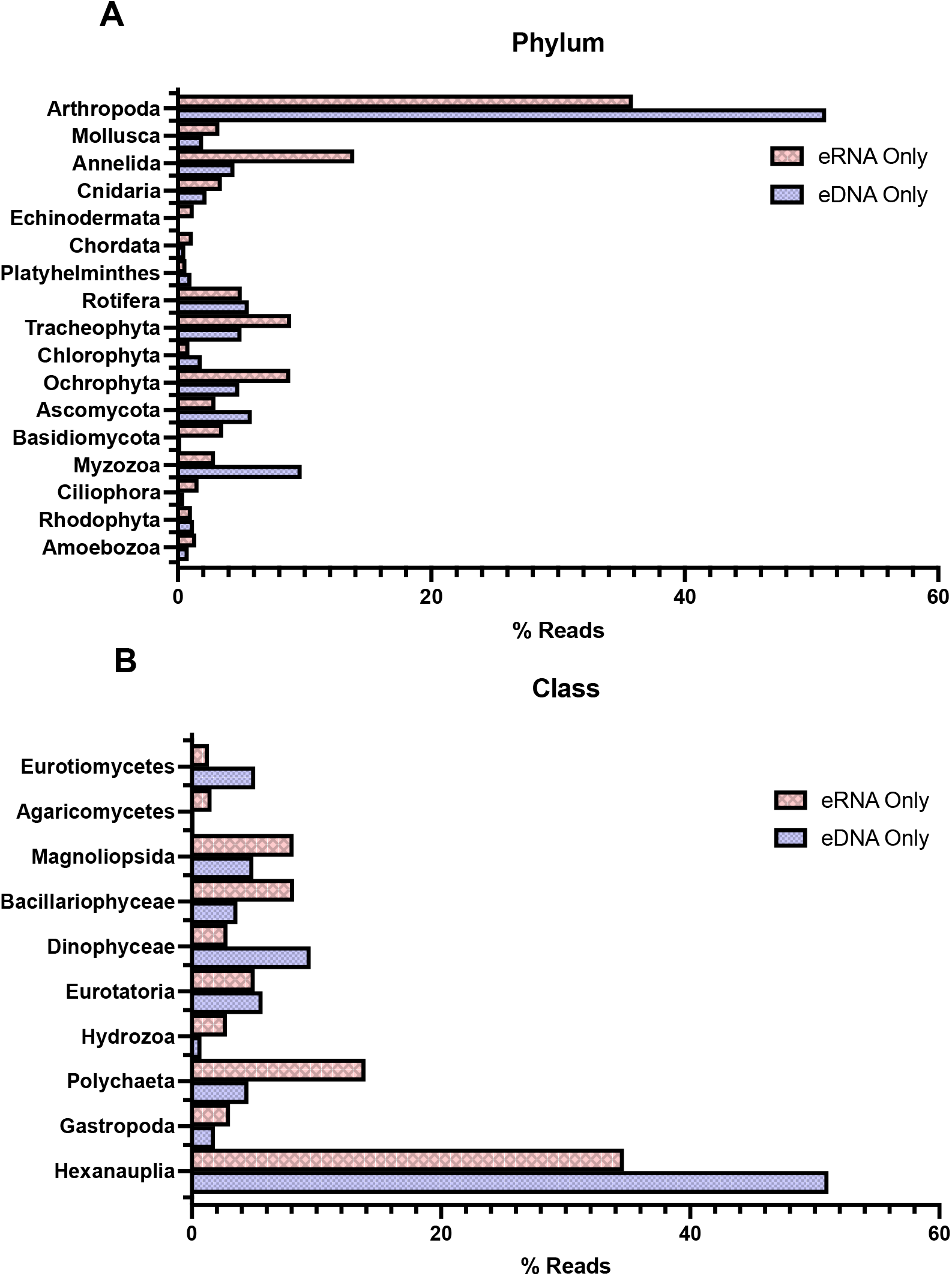
Relative read abundance by (A) phylum and (B) class for eDNA and eRNA.

The two method-unique fractions had broadly similar kingdom profiles, both dominated by Chromista, but differed in magnitude. Of the 624 eRNA-unique OTUs, 317 (50.8%) were Chromista, 147 Animalia, 76 Fungi, 48 Plantae, and 35 Protozoa; of the 305 eDNA-unique OTUs, 141 (46.2%) were Chromista, 95 Animalia, 36 Plantae, 20 Fungi, and 13 Protozoa (Figure 6). To ensure the comparison is not biased toward either template, both unique fractions are shown side by side; this makes clear that the elevated Fungi and Chromista counts in the eRNA-unique set are a difference of degree rather than of kind.

**Figure 6.**
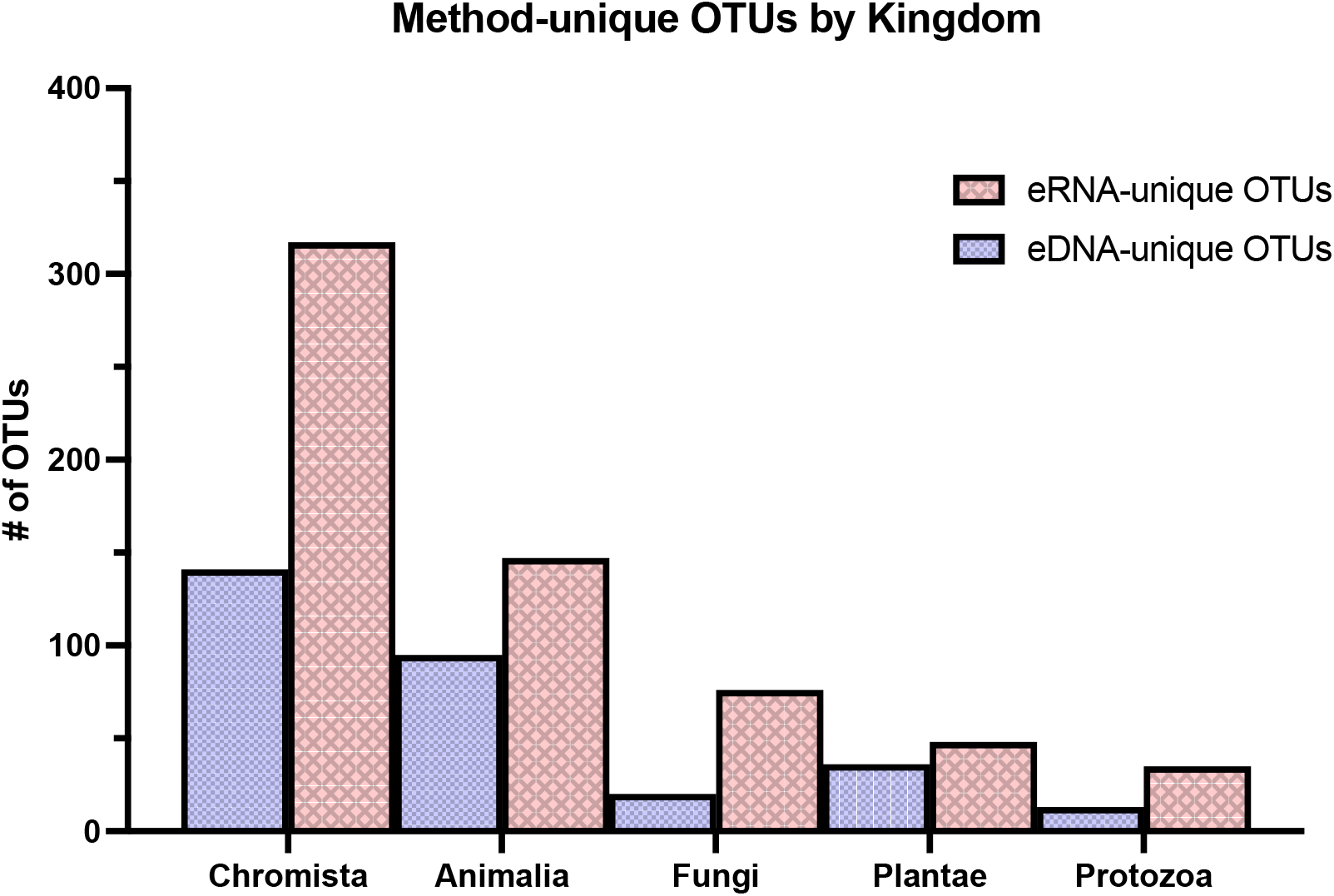
Kingdom composition of the method-unique OTUs, shown for both the eDNA-unique and eRNA-unique fractions.

### 3.7 Community Ordination, PERMANOVA, and PERMDISP

PCoA of Bray-Curtis dissimilarity on the rarefied matrix explained 16.3% and 10.2% of variation on the first two axes (Figure 7). Paired eDNA and eRNA samples from the same site generally fell close together, while habitats such as the cenote, seagrass, and mangrove sites occupied distinct regions of ordination space.

**Figure 7.**
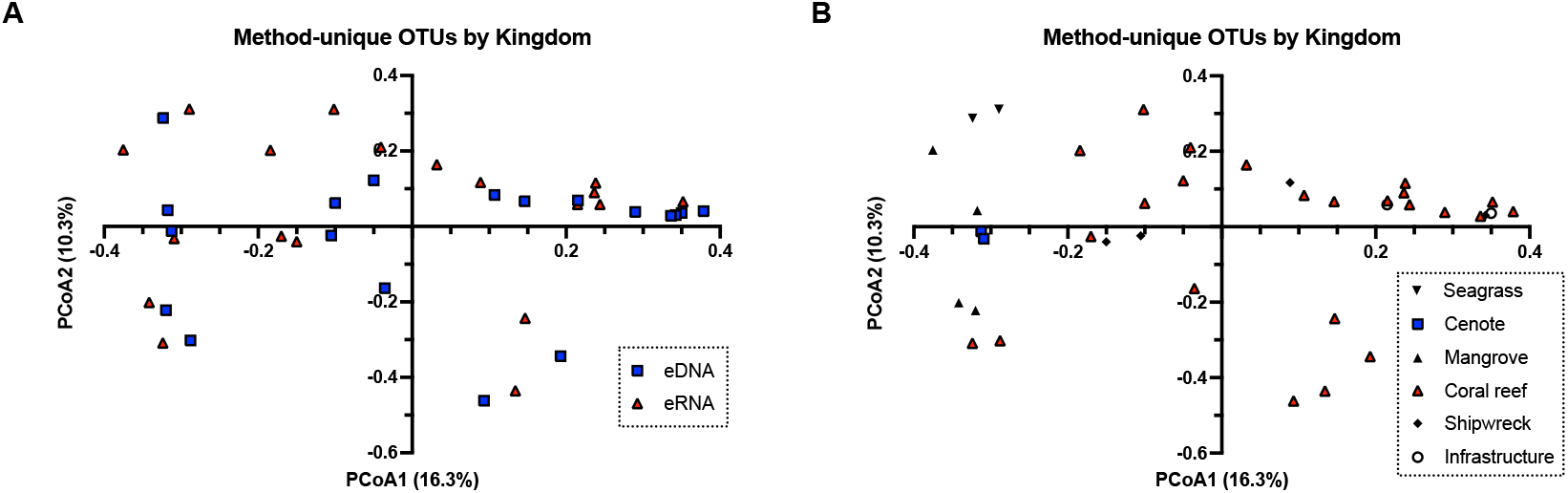
PCoA of Bray-Curtis dissimilarity on rarefied data. (A) Points colored by method, with lines connecting paired eDNA/eRNA samples. (B) Points colored by habitat.

PERMANOVA (Table 3) found that molecular method did not significantly structure abundance-weighted composition (pseudo-F = 1.37, p = 0.107), whereas habitat type did (pseudo-F = 2.489, p = 0.001). Within templates, habitat remained significant for eDNA (F = 1.334, p = 0.036) and was marginal for eRNA (F = 1.251, p = 0.059).

**Table 1.** Site-level OTU richness, shared and method-unique counts, and Jaccard similarity (19 paired sites).

| Site | eDNA | eRNA | Shared | eDNA-only | eRNA-only | Jaccard |
| --- | --- | --- | --- | --- | --- | --- |
| Blue Diamond Shipwreck | 223 | 289 | 127 | 96 | 162 | 0.33 |
| Cenote | 132 | 113 | 81 | 51 | 32 | 0.494 |
| Emmisario Submarino | 207 | 251 | 121 | 86 | 130 | 0.359 |
| Haynes Key | 148 | 191 | 53 | 95 | 138 | 0.185 |
| Johnny Cay | 217 | 274 | 130 | 87 | 144 | 0.36 |
| Mangrove Shallows | 197 | 242 | 158 | 39 | 84 | 0.562 |
| Mangroves - Mural | 90 | 84 | 49 | 41 | 35 | 0.392 |
| Nirvana | 202 | 350 | 182 | 20 | 168 | 0.492 |
| Old Pt Hotel Mar Azul | 173 | 285 | 137 | 36 | 148 | 0.427 |
| Plaza de Toros #1 | 234 | 201 | 104 | 130 | 97 | 0.314 |
| Plaza de Toros #2 | 262 | 273 | 151 | 111 | 122 | 0.393 |
| Punto Marino | 126 | 239 | 113 | 13 | 126 | 0.448 |
| Rocky Cay Shipwreck | 210 | 230 | 112 | 98 | 118 | 0.341 |
| Seaside Reef-Haynes Key | 372 | 410 | 273 | 99 | 137 | 0.536 |
| Shore-side Seagrass | 67 | 89 | 28 | 39 | 61 | 0.219 |
| The Lighthouse #1 | 213 | 332 | 157 | 56 | 175 | 0.405 |
| The Lighthouse #2 | 208 | 196 | 161 | 47 | 35 | 0.663 |
| The Pyramid | 205 | 221 | 126 | 79 | 95 | 0.42 |
| West Point | 330 | 162 | 99 | 231 | 63 | 0.252 |

**Table 2.** Per-site Shannon diversity (H′) for eDNA and eRNA, and which template was higher.

| Site | $H'$ eDNA | $H'$ eRNA | Higher |
| --- | --- | --- | --- |
| Plaza de Toros #1 | 4.035 | 4.325 | eRNA |
| West Point | 4.445 | 4.263 | eDNA |
| Cenote | 2.217 | 1.720 | eDNA |
| Mangrove Shallows | 3.892 | 4.143 | eRNA |
| Haynes Key | 3.384 | 3.082 | eDNA |
| Rocky Cay Shipwreck | 3.586 | 2.794 | eDNA |
| Old Pt Hotel Mar Azul | 3.225 | 3.968 | eRNA |
| Emmisario Submarino | 3.389 | 3.647 | eRNA |
| Blue Diamond Shipwreck | 3.478 | 4.389 | eRNA |
| The Lighthouse #2 | 3.402 | 3.501 | eRNA |
| Plaza de Toros #2 | 2.867 | 3.640 | eRNA |
| Nirvana | 3.288 | 3.948 | eRNA |
| Shore-side Seagrass | 1.110 | 2.222 | eRNA |
| Seaside Reef-Haynes Key | 4.266 | 4.275 | eRNA |
| The Pyramid | 3.091 | 3.247 | eRNA |
| Mangroves - Mural | 2.079 | 2.356 | eRNA |
| Punto Marino | 2.876 | 3.284 | eRNA |
| Johnny Cay | 2.258 | 3.101 | eRNA |
| The Lighthouse #1 | 3.696 | 4.272 | eRNA |

**Table 3.** PERMANOVA and PERMDISP results (999 permutations, Bray-Curtis dissimilarity on rarefied data).

| Test | Samples | Pseudo-F | p-value | Sig. |
| --- | --- | --- | --- | --- |
| Method (eDNA vs. eRNA) | All samples | 1.37 | 0.107 | No |
| Habitat | All samples | 2.49 | 0.001 | Yes |
| Habitat (eDNA only) | eDNA samples | 1.33 | 0.036 | Yes |
| Habitat (eRNA only) | eRNA samples | 1.25 | 0.059 | No |
| PERMDISP: Method | All samples | 0.01 | 0.92 | No |
| PERMDISP: Habitat | All samples | 24.00 | < 0.01 | Yes |

The PERMDISP test clarifies how these results should be read. Between methods, multivariate dispersion was homogeneous (F = 0.014, p = 0.92), so the non-significant method effect reflects a genuine absence of a location difference rather than being masked by unequal dispersion — strengthening the central finding. Among habitats, however, dispersion was significantly heterogeneous (F = 23.995, p < 0.01), which is expected given that several habitats are represented by only one or two sites. The significant habitat PERMANOVA therefore partly reflects differences in within-group spread, and we accordingly treat the habitat effect as indicative rather than definitive.

The apparent tension between the moderate per-site Jaccard similarity (mean 0.40, Section 3.3 and Figure 8) and the non-significant method effect in PERMANOVA is resolved by recognizing that the two metrics measure different things. Jaccard is a presence/absence statistic dominated by rare, low-abundance OTUs, of which each template recovers a different subset; Bray-Curtis used in PERMANOVA is abundance-weighted and dominated by the shared, high-read OTUs (Section 3.4). The methods thus disagree on which rare taxa are present while agreeing on the abundance structure of the community — complementary observations rather than a contradiction.

**Figure 8.**
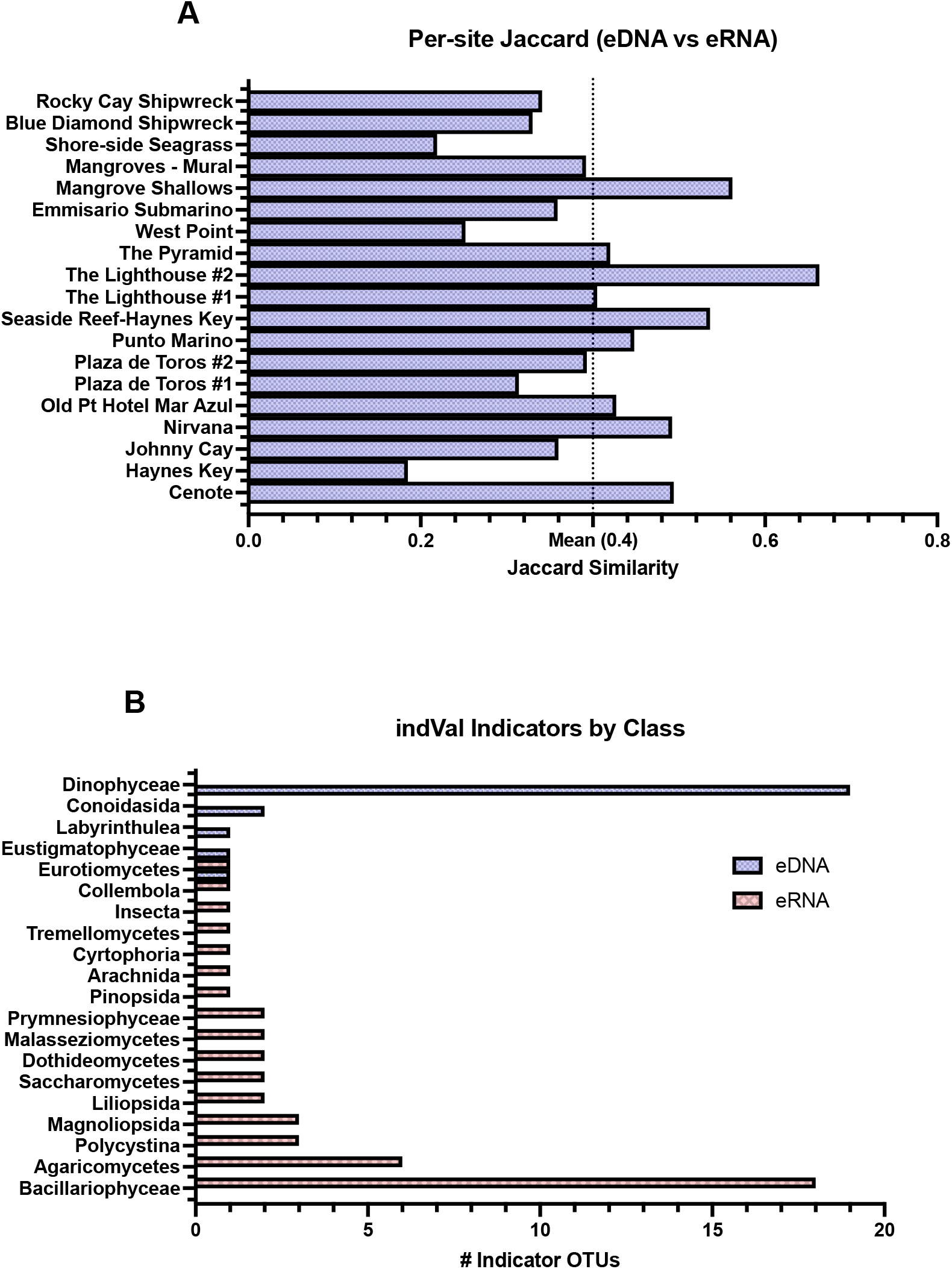
(A) Per-site Jaccard similarity between paired eDNA and eRNA samples (mean = 0.40). (B) Indicator species analysis: number of significant eDNA and eRNA indicator OTUs by taxonomic class.

### 3.8 Indicator Taxa for eDNA and eRNA

Indicator species analysis identified 73 OTUs significantly and preferentially associated with one template (IndVal > 0.3, FDR q < 0.05). The 26 eDNA indicators were dominated by dinoflagellates (Dinophyceae; 19 OTUs), together with the fungus Aspergillus and the dinoflagellate genus Karlodinium. The 47 eRNA indicators were dominated by diatoms (Bacillariophyceae; 18 OTUs) and by fungi (including Agaricomycetes, Saccharomycetes, and Dothideomycetes), with additional terrestrial plant and radiolarian (Polycystina) OTUs.

This contrast provides a community-level, statistically supported basis for the observation that the two templates differ systematically in which microbial-eukaryote groups they emphasize, with eRNA weighted toward diatoms and active fungi and eDNA toward dinoflagellates. We present these as group-level associations rather than claims about individual species’ ecology. A summary by taxonomic class and the full indicator list are provided in the Supplementary Material.

### 3.9 Field-Duplicate Concordance

At the two sites sampled as field duplicates (Plaza de Toros and The Lighthouse), within-method Jaccard similarity (eDNA: 0.316–0.418; eRNA: 0.247–0.285) was of the same order as cross-method Jaccard similarity (0.314–0.663) (Table 4). With only two duplicated sites this is an illustrative observation rather than a formal result, but it suggests that eDNA-versus-eRNA turnover in the rare-taxon fraction is comparable in magnitude to the turnover seen between duplicate samples of the same template.

**Table 4.** Jaccard similarity for within-method duplicate comparisons and cross-method comparisons at the two duplicated sites.

| Comparison | Type | Jaccard |
| --- | --- | --- |
| Plaza de Toros eDNA: sample 1 vs 2 | within-eDNA | 0.316 |
| Plaza de Toros eRNA: sample 1 vs 2 | within-eRNA | 0.247 |
| Plaza de Toros #1: eDNA vs eRNA | cross-method | 0.314 |
| Plaza de Toros #2: eDNA vs eRNA | cross-method | 0.393 |
| The Lighthouse eDNA: sample 1 vs 2 | within-eDNA | 0.418 |
| The Lighthouse eRNA: sample 1 vs 2 | within-eRNA | 0.285 |
| The Lighthouse #1: eDNA vs eRNA | cross-method | 0.405 |
| The Lighthouse #2: eDNA vs eRNA | cross-method | 0.663 |

### 3.10 Illustrative Habitat Case Examples

Beyond the community-level analyses above, several individual sites illustrate the kinds of ecological detail the two templates can surface. We present these as hypothesis-generating case examples, not as quantitatively supported food-web reconstructions: most involve single unreplicated sites, and co-detection of taxa in a water sample does not by itself establish a trophic linkage.

At the Rocky Cay shipwreck, several biofouling-associated taxa were far more abundant in eRNA than eDNA, including the hydrozoan *Clytia hemisphaerica* (23,707 vs. 280 reads) and the polychaete *Scolelepis* sp. (72,584 vs. 1,143 reads), with *Branchiomma* sp. recovered almost entirely in eRNA. In the mangrove sites, eRNA added benthic taxa such as the copepod *Longipedia koreana* (2,246 vs. 883 reads) to the planktonic signal seen in eDNA; we note that the difference for the polychaete *Syllis pigmentata* (891 vs. 462 reads) is based on two data points and is not presented as a meaningful trend. At the Shore-side Seagrass site, eDNA was dominated by terrestrial plant material (Solanaceae; roughly 78% of reads), whereas eRNA recovered a more marine-weighted assemblage including copepods and dinoflagellates — the clearest case in which eRNA reduced masking by terrestrial input. On coral reefs, the appendicularian *Fritillaria borealis* (1,016 vs. 289 reads) and *Symbiodiniaceae*/cnidarian signals illustrate taxa recovered by both templates; with single sites these remain anecdotal.

The potentially management-relevant detections in the dataset — for example the ciguatera-associated dinoflagellate *Gambierdiscus caribaeus*, recovered at six sites — are likewise reported as occurrence observations that would require targeted follow-up before any inference about risk.

## 4. Discussion

### 4.1 eRNA as a Complementary Layer

Across 19 paired sites, eRNA recovered significantly higher OTU richness and Shannon diversity than eDNA and contributed 624 unique OTUs (32.1% of the total). The read-weighted analysis shows these unique detections are overwhelmingly rare taxa, so eRNA’s contribution is best framed as extending the detected tail of the community — particularly metabolically active microbial eukaryotes and some benthic invertebrates — rather than as overturning the picture of dominant community structure that eDNA already provides. We are careful not to over-state this as a wholesale difference in community composition, because the abundance-weighted ordination shows the two templates largely agree (Section 4.2).

### 4.2 Method Versus Habitat as Drivers of Composition

Our central multivariate result is that habitat type, not molecular method, structures abundance-weighted community composition. By “genuine ecological signal” we mean specifically this: the variation attributable to habitat (a biological property of the sites) exceeds and is statistically separable from the variation attributable to the choice of template (a methodological property). The non-significant method effect, combined with homogeneous between-method dispersion in PERMDISP, indicates that an eDNA and an eRNA sample from the same site resemble each other about as much as expected for the same community assayed two ways.

We temper the habitat side of this comparison. PERMDISP found significantly heterogeneous dispersion among habitats, which is unsurprising given that several habitats were represented by only one or two sites. Part of the significant habitat PERMANOVA therefore reflects unequal within-group spread rather than pure location differences, and the habitat effect should be read as indicative. The well-replicated comparison in this study is the method comparison; the habitat comparison is exploratory and would benefit from balanced replication across habitats.

### 4.3 Reconciling Presence/Absence and Abundance-Weighted Views

The moderate mean Jaccard similarity (0.40) and the non-significant PERMANOVA method effect are sometimes read as contradictory. They are not: Jaccard is presence/absence and is governed by the many rare OTUs that each template samples differently, whereas Bray-Curtis is abundance-weighted and governed by the shared high-abundance OTUs. The consistent message across both is that the templates diverge in the rare-taxon tail and converge in the abundant core.

### 4.4 Indicator Taxa and the Activity Hypothesis

The indicator analysis gives the clearest mechanistic hint in the dataset: eRNA indicators are dominated by diatoms and active fungi and eDNA indicators by dinoflagellates. This is consistent with the hypothesis that eRNA is weighted toward metabolically active cells with high ribosomal content, although marker-specific amplification efficiency and database completeness are alternative contributors we cannot fully exclude with a single COI marker. We therefore present the indicator contrast as a robust descriptive pattern with a plausible activity-based interpretation, not as proof of differential activity.

### 4.5 Rarefaction and Robustness

Rarefaction to the minimum observed depth did not change the richness conclusions, supporting the view that the eRNA-richness advantage is not a sequencing-effort artifact. We frame this as a supporting robustness check; given ongoing debate over rarefying versus model-based normalization for metabarcoding data, we avoid resting any conclusion solely on a single normalization choice.

### 4.6 Read-Weighted Context for Monitoring

For monitoring that targets dominant community composition, eDNA alone captures most of the abundance-weighted signal. For comprehensive inventories, rare-taxon detection, or indicator-based surveillance, adding eRNA materially expands coverage. The cost of the additional RNA workflow should be weighed against these specific objectives.

### 4.7 Terrestrial-Input Masking: The Clearest Use Case

The strongest, most defensible use case in our data is the seagrass site, where eDNA was dominated by terrestrial plant material and eRNA recovered a more marine assemblage. In systems with strong terrestrial inputs — estuaries, mangrove fringes, run-off-influenced coasts — the faster turnover of RNA may reduce masking by allochthonous DNA, improving detection of the resident marine community. This is a concrete, mechanistically plausible advantage that warrants targeted, replicated testing.

### 4.8 Limitations

Several limitations bound our conclusions. (1) Most sites were sampled once per template, and only two sites were field-duplicated; the case examples in Section 3.10 are unreplicated and hypothesis-generating. (2) Habitat replication was strongly unbalanced (coral reef n = 12 versus single sites for several habitats), and PERMDISP confirmed heterogeneous habitat dispersion, so habitat effects are indicative rather than definitive. (3) A single COI marker limits phylogenetic breadth and is subject to primer bias; multi-marker panels would broaden coverage. (4) Co-detection of taxa does not establish trophic interaction. (5) The RNA stabilization and reverse-transcription workflow introduces uncertainties (incomplete DNase removal, RT efficiency) that paired no-RT controls mitigate but do not eliminate. (6) Environmental covariates were not measured alongside sampling, limiting interpretation of habitat effects.

### 4.9 Potential Health and Management Indicators

The dataset contained taxa of potential management interest, including the ciguatera-associated dinoflagellate *Gambierdiscus caribaeus* at six sites, elevated *Aspergillus* at two sites, and molecular signatures consistent with land-to-sea input. We report these as occurrence-level observations to flag for targeted follow-up; the present design does not support quantitative risk inference.

### 4.10 Toward a Monitoring Framework

Rather than prescribing paired eDNA/eRNA as a universal standard, we suggest it be considered where its specific advantages apply: settings with strong terrestrial masking, programs needing rare-taxon or indicator surveillance, and studies of community activity. Where these do not apply, eDNA alone may remain the most cost-effective choice. We further suggest that future paired studies adopt balanced habitat replication, biological (not only field) replicates, and — where feasible — multi-marker panels and paired environmental covariates.

## 5. Conclusions

In a paired eDNA/eRNA survey across 19 tropical marine sites, eRNA recovered significantly higher OTU richness and Shannon diversity than eDNA and contributed a substantial set of unique, predominantly rare OTUs enriched for diatoms and fungi. Abundance-weighted community composition was structured by habitat rather than by molecular method, and PERMDISP confirmed that the absence of a method effect was not an artifact of unequal dispersion. Indicator analysis showed a systematic contrast — diatoms and active fungi associated with eRNA, dinoflagellates with eDNA — consistent with, though not proof of, an activity bias in eRNA. The clearest practical advantage of eRNA in our data was reduced masking by terrestrial DNA at the seagrass site. We offer these results as evidence that eRNA is a useful complementary layer for marine biodiversity assessment in specific contexts, and as a foundation — building on prior mesocosm and freshwater work — for more fully replicated paired designs.

## Author Contributions

Author contributions are described below following the CRediT (Contributor Roles Taxonomy) framework. S. Bedingfield performed the field sampling and sample collection, with the coordination and support of C. Vanegas Moreno and the scientific staff of CORALINA. A. More and S. Bedingfield developed the original study design and conceptualization. [PLACEHOLDER text below should be completed once the final author list and individual roles are confirmed. More to be done.]

## Acknowledgments

We gratefully acknowledge Carolina Vanegas Moreno and the scientific staff of CORALINA (the Corporation for the Sustainable Development of the Archipelago of San Andrés, Providencia and Santa Catalina), the regional environmental authority responsible for managing the natural resources and environment of the archipelago and the Seaflower Biosphere Reserve, for their coordination and support of the field sampling. We thank Maria Jose Abuabara for providing and supporting the initial concept and for establishing the partnerships that made this work possible. We thank EQO for sample collection equipment support and eRNA/eDNA extraction from the sample membranes and NatureMetrics for subesequent laboratory processing, sequencing, and bioinformatic analysis. We also thank the communities of San Andrés for their support during fieldwork.

## Data Availability

The data supporting the findings of this study are described in the main text and Supplementary Materials. Raw data and processed OTUs can be accessed here: https://doi.org/10.5281/zenodo.20535111

