## Supplemental Materials for "Complementary Insights from Environmental DNA and Environmental RNA Metabarcoding for Marine Biodiversity Assessment Around San Andrés Island, Colombia"

### Supplementary Material

#### S1. Data Provenance and the Duplicated eRNA Block

The OTU table supplied for this project contained two blocks of eRNA columns. One block (19 samples) is internally consistent and reproduces every site-level statistic reported here. The second block (a parallel set of 19 columns) contained thousands of negative values (minimum below −45,000) that are not interpretable as read counts. All analyses in this manuscript use the consistent block only; the second block was excluded as non-canonical.

#### S2. Legacy 20-Site Overlap

If the eDNA-only Old Point site is included, the all-sites totals are 1,961 OTUs (1,342 eDNA, 1,639 eRNA; 1,020 shared, 322 eDNA-unique, 619 eRNA-unique). The paired 19-site totals used throughout the main text are 1,944 / 1,320 / 1,639 / 1,015 / 305 / 624. The two differ only in the unpaired site and no conclusion depends on the choice; the paired totals are used in the main text so that the eDNA-versus-eRNA comparison rests only on sites where both templates were obtained.

#### S3. Observed Versus Rarefied Richness

Rarefaction to 16,758 reads changed observed richness only marginally at every site, and the rank order of sites and the eRNA-higher-than-eDNA pattern (14/19) were preserved. The full observed-versus-rarefied table and per-sample rarefaction curves are provided as supplementary files (see the accompanying GraphPad data workbook, sheets S_Rarefied and F3_Richness).

#### S4. Combined-Diversity Computation

A pooled-dataset Shannon diversity can be computed by summing eDNA and eRNA reads per OTU and recomputing H′ on the pooled counts. When two templates differ strongly in evenness, this pooling can lower H′ relative to the more even template, so a pooled value does not behave as a simple sum of information. Because this is a property of pooling rather than a biological result, pooled diversity is documented here for completeness rather than used as a main finding.

#### S5. Indicator Species — Full Results

The complete list of the 73 significant indicator OTUs (IndVal > 0.3, FDR $q < 0.05$), with taxonomy, indicator value, and adjusted p-value, is provided in the accompanying data workbook (sheet F7_IndVal).

#### S6. Data Anomalies Log

A consolidated log of data-provenance and quality-control items — the duplicated eRNA column block, the paired-versus-all-sites overlap totals, the per-site Shannon comparison (eRNA higher at 15 of 19 sites), the DNA-quantification assay, and the site-coordinate corrections described in S7 — is provided in the data workbook (sheet DATA_ANOMALY) so that all items are tracked in one place.

#### S7. Site Coordinates

Coordinates below are taken from the field sample manifest. Two longitudes were recorded as positive values (Plaza de Toros #2 and Punto Marino); these were set to the western hemisphere (negative) to match all other sites and the known position of the island, which places both sites in their expected locations relative to the coast. The field manifest also notes that the site labeled “Punto Marino” may correspond to a location locally known as “Mundo Marino”; the label “Punto Marino” is retained throughout for consistency with the sequence data identifiers.

**Table S1. Site coordinates (decimal degrees) and habitat, ordered north to south.**

| Site | Latitude (°N) | Longitude (°W) | Habitat |
|---|---|---|---|
| Johnny Cay | 12.60060 | -81.69218 | Coral reef |
| Plaza de Toros #2 | 12.58514 | -81.71908 | Coral reef |
| Plaza de Toros #1 | 12.58514 | -81.71908 | Coral reef |
| The Pyramid | 12.58435 | -81.68349 | Coral reef |
| Cenote | 12.58040 | -81.71410 | Cenote |
| Mangrove Shallows | 12.57798 | -81.69864 | Mangrove |
| Emmisario Submarino | 12.57209 | -81.72634 | Infrastructure |
| Mangroves - Mural | 12.57033 | -81.70716 | Mangrove |
| Mangrove - Old Point | 12.56350 | -81.70727 | Mangrove (eDNA only) |
| Old Pt Hotel Mar Azul | 12.55837 | -81.70177 | Coral reef |
| Punto Marino | 12.55553 | -81.69626 | Coral reef |
| Seaside Reef-Haynes Key | 12.55368 | -81.67889 | Coral reef |
| Haynes Key | 12.55091 | -81.69109 | Coral reef |
| Shore-side Seagrass | 12.54612 | -81.70348 | Seagrass |
| Rocky Cay Shipwreck | 12.54115 | -81.70081 | Shipwreck |
| West Point | 12.52090 | -81.73119 | Coral reef |
| Blue Diamond Shipwreck | 12.51780 | -81.73168 | Shipwreck |
| The Lighthouse #1 | 12.51625 | -81.73080 | Coral reef |
| The Lighthouse #2 | 12.51549 | -81.73053 | Coral reef |
| Nirvana | 12.50128 | -81.73213 | Coral reef |
